# Uropathogenic *E. coli* induces DNA damage in the bladder

**DOI:** 10.1101/2020.05.07.080291

**Authors:** Camille V. Chagneau, Clémence Massip, Nadège Bossuet-Greif, Christophe Fremez, Jean-Paul Motta, Ayaka Shima, Céline Besson, Pauline Le Faouder, Nicolas Cénac, Marie-Paule Roth, Hélène Coppin, Maxime Fontanié, Patricia Martin, Jean-Philippe Nougayrède, Eric Oswald

## Abstract

Urinary tract infections (UTIs) are among the most common outpatient infections, with a lifetime incidence of around 60% in women. We analysed urine samples from 223 patients with community-acquired UTIs and report the presence of a metabolite released during the synthesis of colibactin, a bacterial genotoxin, in 50 of the samples examined. Uropathogenic *Escherichia coli* strains isolated from these patients, as well as the archetypal *E. coli* strain UTI89, were found to produce colibactin. In a murine model of UTI, the machinery producing colibactin was expressed during the early hours of the infection, when intracellular bacterial communities form. We observed extensive DNA damage both in umbrella and bladder progenitor cells. To the best of our knowledge this is the first report of colibactin production in UTIs in humans and its genotoxicity in bladder cells. This bacterial genotoxin, which is increasingly suspected to promote colorectal cancer, should also be scrutinised in the context of bladder cancer.

## Introduction

Urinary tract infections (UTIs) are one of the most common bacterial infections, affecting approximately 150 million individuals each year ^1^. UTIs occur most frequently in women, with more than 60% of females diagnosed with a UTI during their lifetime ^2^. The severity of these infections ranges from asymptomatic bacteriuria and cystitis, *i.e.* infections localised to the bladder, to urosepsis, which can be fatal. Recurrences are very frequent, since approximately 30% of women experience a new UTI episode after resolution of the initial infection ^2^. In addition to their consequences in terms of morbidity, mortality and associated economic and societal losses, UTIs are also a major reason for antibiotic treatments and thus strongly contribute to the global issue of antibiotic resistance. *Escherichia coli* strains, termed uropathogenic *E. coli* (UPEC) cause approximately 80% of all UTIs. These strains belong mainly to phylogroup B2, which is increasingly present in the intestinal microbiota, the reservoir of UPEC ^3,4^. UPEC strains produce a large number of virulence factors ^5–7^. In particular, several toxins have long been associated with UPEC pathogenicity, such as α-hemolysin and CNF1 toxins ^8,9^. More recently, a large proportion of UPEC strains which carry *pks* pathogenicity island encoding the genotoxin colibactin have been described ^10–13^.

The *pks* pathogenicity island, composed of *clbA-S* genes, encodes a polyketide-non-ribosomal-peptide (PK-NRP) biosynthesis machinery ^14^. Colibactin is first synthesised as an inactive prodrug by the sequential interventions of Clb enzymes. ClbP peptidase subsequently cleaves the C14-Asparagine (C14-Asn) motif thereby releasing the mature, active form of colibactin with its twin warheads (Fig. 1a) ^15–17^. The genotoxin alkylates adenine residues on both strands of DNA, producing DNA interstrand cross-links ^18–20^. These highly toxic DNA lesions initiate a DNA damage response, by phosphorylating replication protein A (pRPA) and phosphorylating the H2AX histone variant (pH2AX) (Fig. 1b) ^14,18^. Incomplete repair of this DNA damage can result in gene mutations ^21^. *E. coli* strains carrying *pks* island have been shown to promote colon carcinogenesis in different mouse models ^22–24^. In epidemiological studies, *pk*s+ *E. coli* strains are more prevalent in the gut microbiota of patients with colorectal cancer and a distinct mutational signature in human cancer genomes, predominantly colorectal tumours, was recently associated with colibactin genotoxic activity, further implicating an involvement of colibactin-producing *E. coli* in tumorigenesis ^22,23,25,26^. This mutational signature has also been identified in tumours of the urinary tract ^26^.

**Fig. 1:**
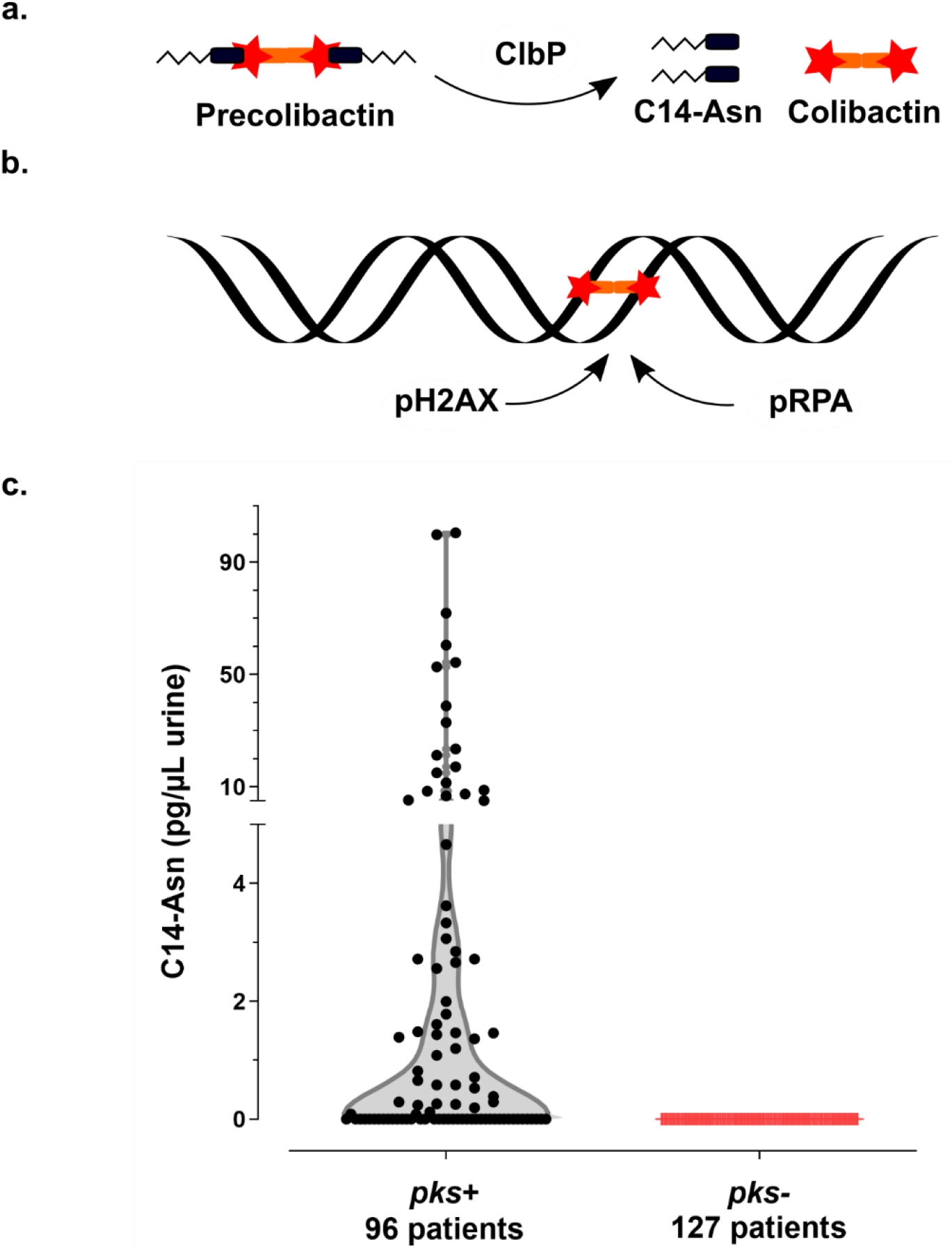
Synthesis of the genotoxin, colibactin, releases the C14-Asn metabolite detected in the urine of patients with UTI. **a.** Colibactin is first synthesised as a prodrug, precolibactin and then cleaved by ClbP peptidase thereby releasing the mature genotoxic dimeric colibactin, and C14-Asn. **b.** Colibactin alkylates both strands of the DNA helix, generating an interstrand cross-link. In response to this DNA damage, the host cell machinery recruits and phosphorylates the H2AX and RPA proteins. **c.** Concentration of C14-Asn in human urines according to the presence of a *pks* island in the genome of the corresponding UPEC isolate as determined by LC-MS/MS. All individual data points are shown on a violin plot, with a two segments linear Y range. C14-Asn was below the LC-MS/MS detection limit in the urine of patients infected with UPEC isolates that did not contain a *pks* island.

Our current study shows that colibactin producing bacteria induce DNA damage in bladder cells, including in urothelial regenerative cells and that colibactin is produced by *pks*+ UPEC clinical strains isolated from human UTIs.

## Results

### Evidence of colibactin production in the urine of patients infected with UPEC

We collected urine samples from 223 adult patients with community-acquired pyelonephritis, cystitis or asymptomatic bacteriuria caused by *E. coli* at the University Hospital of Toulouse, France. Urine samples were analysed for the presence of C14-Asn, the aminolipid released during the final colibactin maturation step (Fig. 1a). In contrast to the highly reactive and unstable colibactin, C14-Asn is stable and can be quantified by LC-MS/MS. C14-Asn was detected in urine samples of one quarter (55/223) of UTI patients, including asymptomatic infections (Fig. 1c & Table S1). We isolated *E. coli* strains harbouring genomic *pks* island from all urine samples which also contained C14-Asn. Conversely, C14-Asn was below the LC-MS/MS detection limits in urine samples of patients infected with *E. coli* strains which did not carry the *pks* pathogenicity island.

### Phylogenetic distribution of *pks* island in UPEC strains

All 223 *E. coli* strains isolated during this sampling campaign were whole genome sequenced. The phylogenetic distribution of these *E. coli* isolates was typical of strains which cause UTIs ^2^. A majority of strains belonged to the phylogroups B2 (69%) and D (15%) (Fig. 2 & Table S1). Forty three percent of strains harboured *pks* island. All the *pks*+ strains belonged to the phylogenetic group B2 and to the most common sequence types (STs) of extra-intestinal pathogenic *E. coli* such as ST73, ST95, ST141 and ST404 (Fig. 2). As expected for extra-intestinal pathogenic *E. coli*, multiple known or suspected virulence genes were also present in the genome of these *pks*+ strains. *pks* island are thus widely present in the typical UPEC strains that cause community-acquired UTIs.

**Fig. 2:**
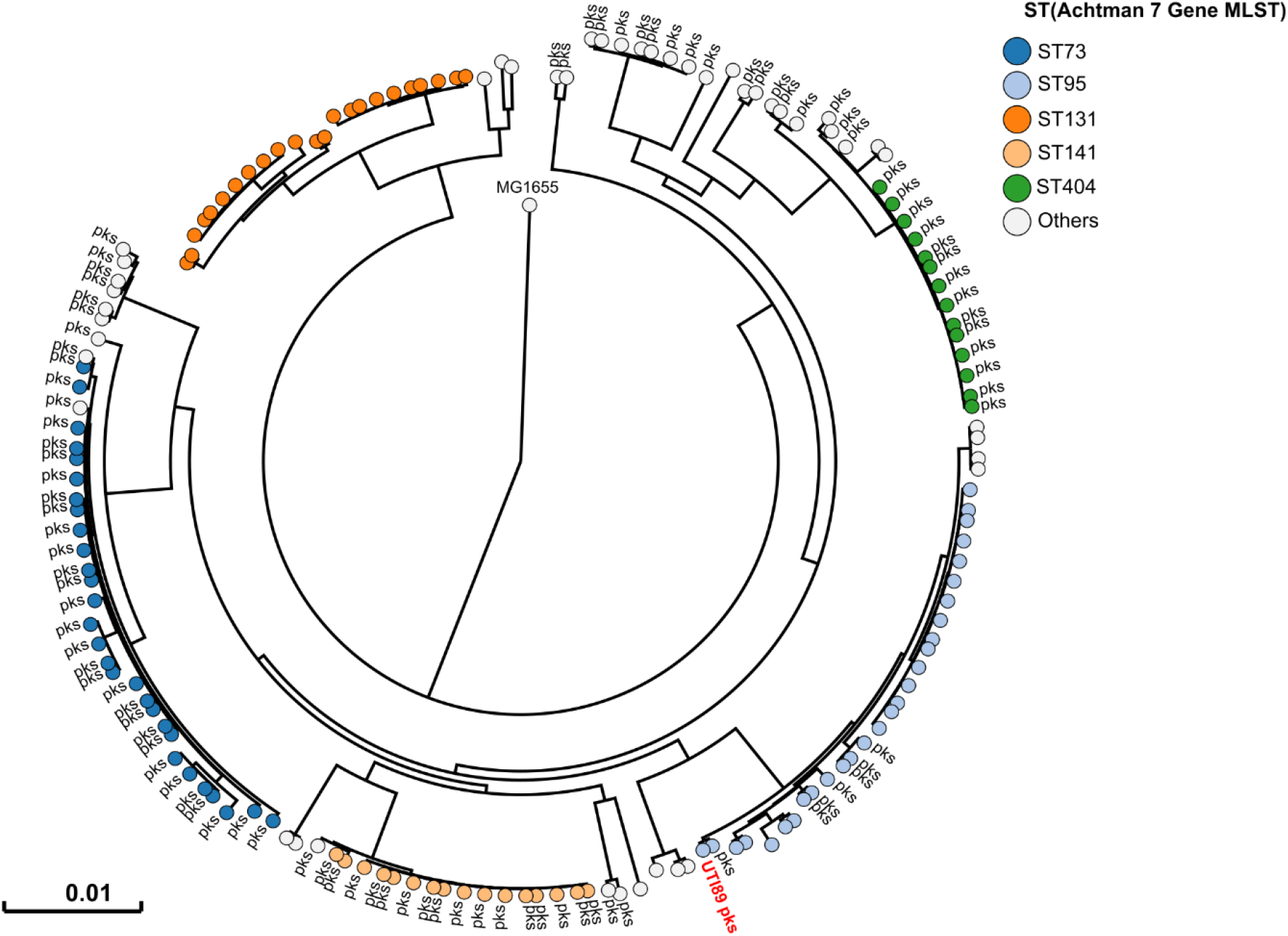
Human *pks*+ UPEC belong to major lineages of extraintestinal pathogenic *E. coli* from phylogroup B2. A phylogenetic tree based on whole genome analysis was constructed and rooted on *E. coli* MG1655. Each circle represents an individual UPEC strain. Main Sequence Types are grouped by colours. The presence of a *pks* island is denoted adjacent to each circle. The archetypal UPEC strain UTI89, which is *pks*+, was also included in this tree.

### UPEC strains carrying *pks* island produce the genotoxin colibactin

We next sought to confirm that *pks*+ UPEC strains expressed colibactin. We first evaluated the ability of UPEC strain UTI89, commonly used in rodent models of UTI, to produce colibactin and to be genotoxic. We detected the colibactin C14-Asn cleavage product in bacterial cultures (Fig. 3a) and confirmed the strain’s genotoxicity on exogenous double-stranded DNA (Fig. 3b). These effects were comparable to those observed with strain NC101 (Fig. 3a-b) which has been shown to be pro-carcinogenic in different colorectal cancer mouse models. The same genotoxic effects were also observed with three clinical *pks+* UPEC isolates from asymptomatic bacteriuria, cystitis or pyelonephritis cases (Fig. 3b). As was shown for NC101, infecting human epithelial cells with *pks+* UPEC strains induced the formation of nuclear pRPA and pH2AX foci, indicating that the DNA of exposed cell is damaged (Fig. 1b, Fig. 3c). All together, these results demonstrate that UPEC strains *pks* island are functional, mediate the synthesis of colibactin and are genotoxic.

**Fig. 3:**
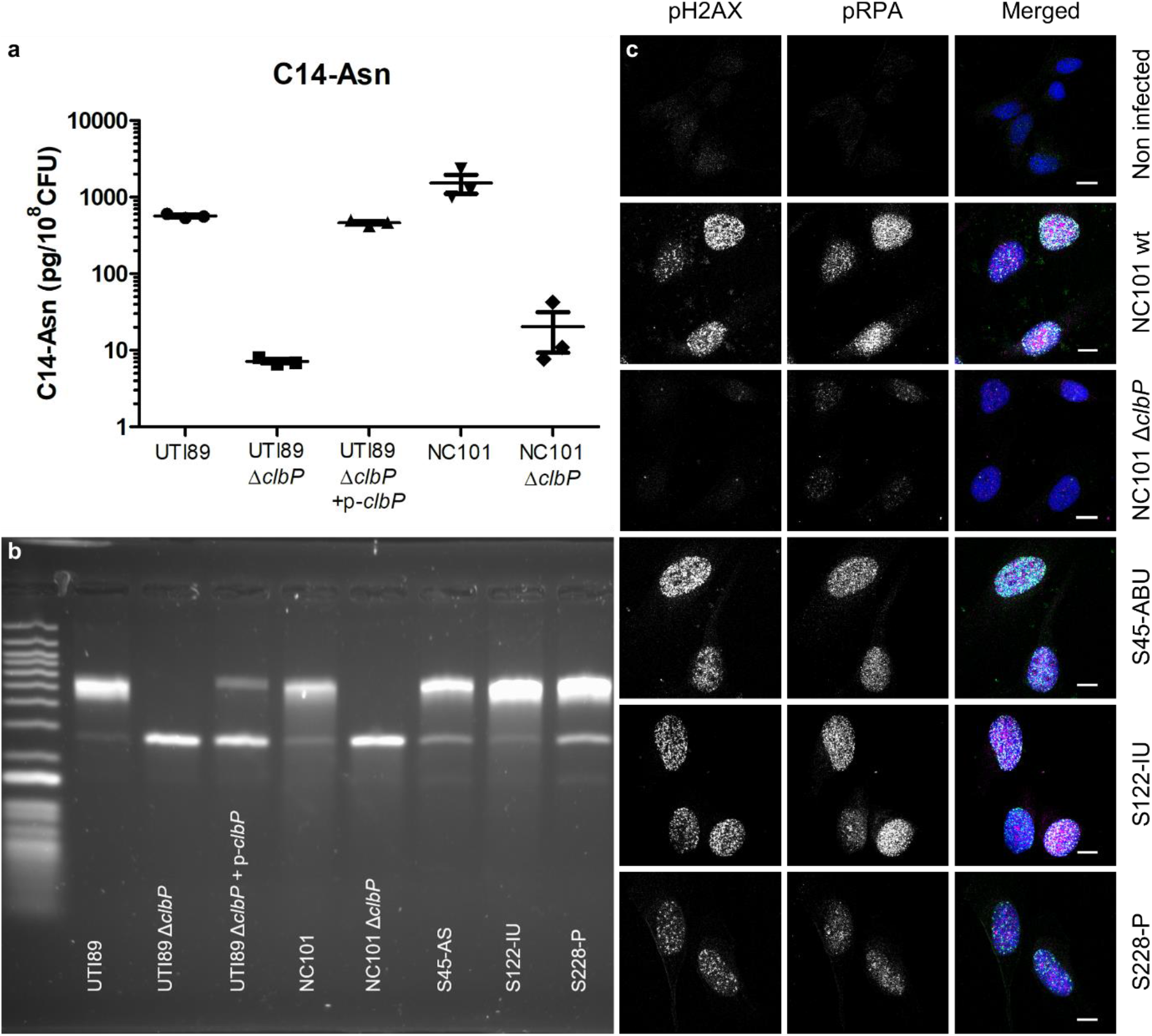
The archetypal cystitis *E. coli* strain UTI89 and clinical *pks*+ UPEC isolates produce colibactin and induce DNA damage. **a**. LC-MS/MS quantification of the C14-Asn cleavage product released following colibactin maturation by ClbP peptidase in bacterial pellets of *E. coli* strains UTI89 and NC101 and their respective Δ*clbP* mutants. Quantifications were performed in triplicate and represented as a mean ± standard error of the mean (SEM). **b.** DNA interstrand cross-links formed after exposure of linear double stranded DNA to UTI89, NC101 and their respective Δ*clbP* mutants as well as three representative clinical human UPEC isolates, visualised by electrophoresis under denaturing conditions. DNA with interstrand cross-links migrates at an apparent molecular weight of twice that of DNA without cross-links. In the absence of any crosslinking DNA migrates as a single denatured strand. S45, S122, S228: Human UPEC strains from AS=asymptomatic bacteriuria; IU=cystitis; P=pyelonephritis respectively. **c.** S33p-RPA32 and pH2AX immunofluorescence staining of HeLa cells, 16 hours after infection with the reference NC101 strain, its Δ*clbP* mutant or human UPEC strains (S45, S122, S228). pH2AX and S33p-RPA32=pRPA: grayscale. Merged: green=pH2AX; magenta=S33p-RPA32; blue=DAPI; scale bar=10 μm.

### Colibactin is produced during UTIs and induces bladder urothelium DNA damage

In order to monitor the expression of the *pks* synthesis machinery during a UTI, we transformed UPEC strain UTI89 with a plasmid expressing a GFP tagged *pks* island-encoded polyketide ClbI synthase from its endogenous promoter. In bladder tissue collected 6 hours after infection, we observed ClbI-GFP expressing bacteria in intracellular bacterial communities (IBCs) inside superficial umbrella cells which line the lumen of the bladder (Fig. 4a-d & S1). Production of colibactin was further confirmed by the detection of C14-Asn in pooled urines from 8 mice 24 hours after infection (C=0.66 pg/μL). We next assessed whether the metabolically active *pks* machinery was associated with DNA damage in bladder cells. Phosphorylation of H2AX was readily detected in nuclei of umbrella cells containing IBCs, 6 hours after infection with wild-type *E. coli* UTI89 with or without the p-*clbI*-*gfp* plasmid (Fig. 4c-d & S2). A *clbP* mutant unable to produce colibactin was very weakly genotoxic or not genotoxic at all (Fig. S2), although equally capable of colonising the bladder (Fig. 4l) and inducing the formation of IBCs (Fig. 4e-k). These results show that colibactin is expressed *in vivo,* in the bladder from the very early stages of UTI and can induce DNA damage.

**Fig. 4:**
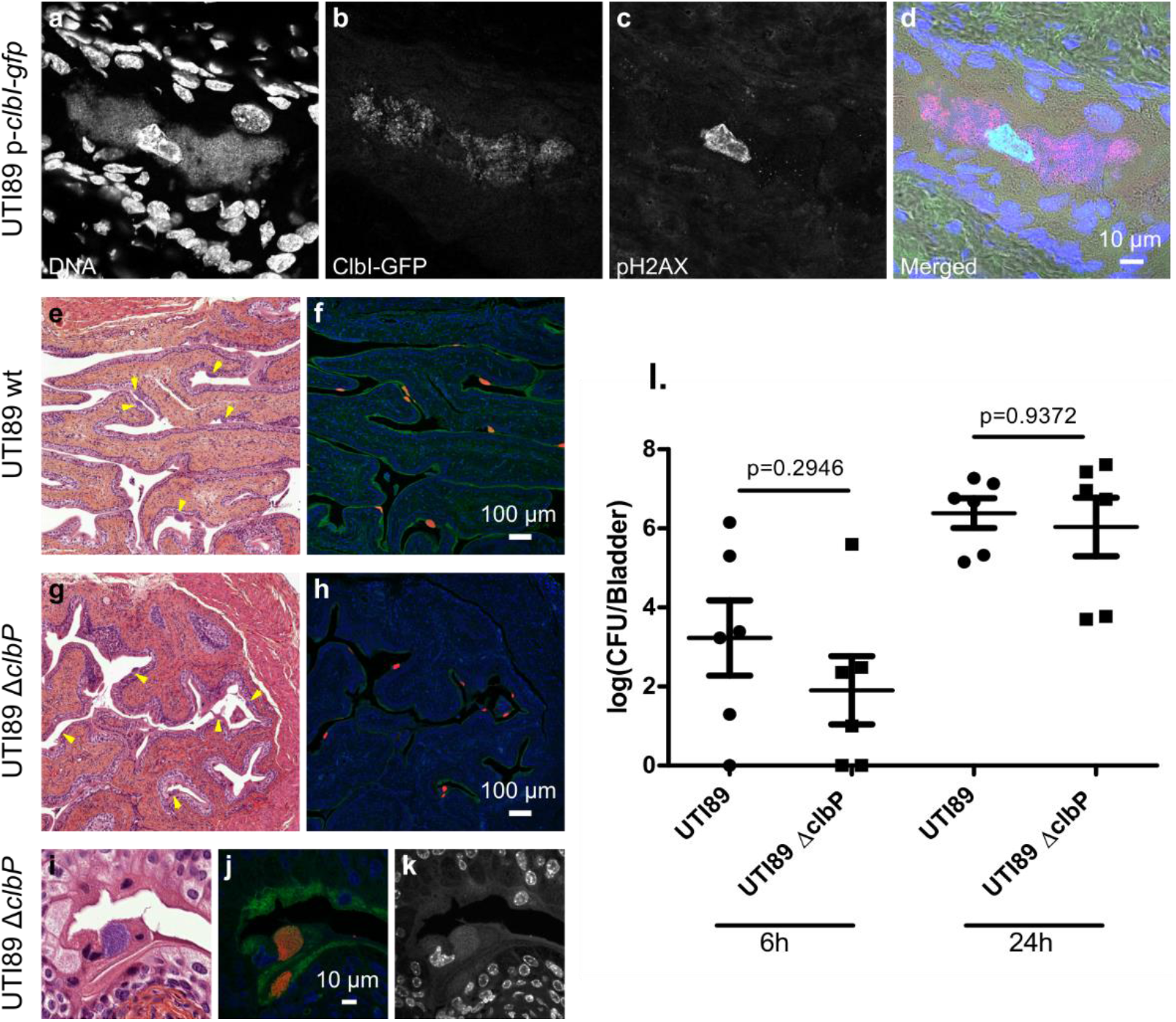
Colibactin is produced during UTIs and induces DNA damage. **a-d**. Confocal microscopic detection of ClbI-GFP expression and pH2AX in frozen bladder sections containing an IBC, 6 hours post infection with UTI89, hosting the p-*clbI*-*gfp* fusion plasmid. The individual channel images are shown in grayscale. In the merged (d) image: blue=DNA, magenta=ClbI-GFP, green=pH2AX, grey=phase contrast. **e-k.** Bladder sections 6 hours post inoculation with wild-type UTI89 (e, f) or the isogenic Δ*clbP* mutant (g-k) stained with haematoxylin-eosin (e, g, i), FISH (f, h, j) or DAPI (k, grayscale) all of which detect IBCs (arrows). In the FISH images: blue=DAPI stain DNA; green= FITC-conjugated wheat germ agglutinin (WGA) stained polysaccharides; red=bacterial cells, labelled with the universal 16S FISH probe. **l.** Bladder tissue *E. coli* counts, 6 h and 24 h after infection with wild-type *E. coli* UTI89 (circles) or Δ*clbP* (squares). Each data point corresponds to one mouse, with mean ± standard error of the mean (SEM) shown for each group. n=6. Mann-Whitney U test.

### Colibactin induces urothelial cell DNA damage in the regenerative compartment

Bladders from mice infected with wild-type *E. coli* UTI89 exhibited pH2AX positive nuclei in superficial umbrella cells, but also in urothelial basal cells (Fig. 5a-c). Bladders from mice infected with the *clbP* mutant did not exhibit any pH2AX positive cells. pH2AX positive cells nevertheless reappeared after complementation with the wild-type *clbP* allele (Fig. 5d-f & S3). The basal urothelium compartment harbours keratin-14 positive (Krt14+) progenitor cells, which are important for urothelium renewal following injury. There was an overrepresentation of Krt14+ cells in urothelial tissue infected with wild-type or with *clbP* mutant UTI89 (Fig. S4). Importantly, bladders infected with the genotoxic wild-type UTI89 strain exhibited basal urothelial cells that were positive for both the regenerative cell marker Krt14 and the DNA damage marker pH2AX (Fig. 5g-l). Thus, during UTI with a *pks+* UPEC, colibactin induces DNA damage in superficial and basal regenerative urothelial cells.

**Fig. 5:**
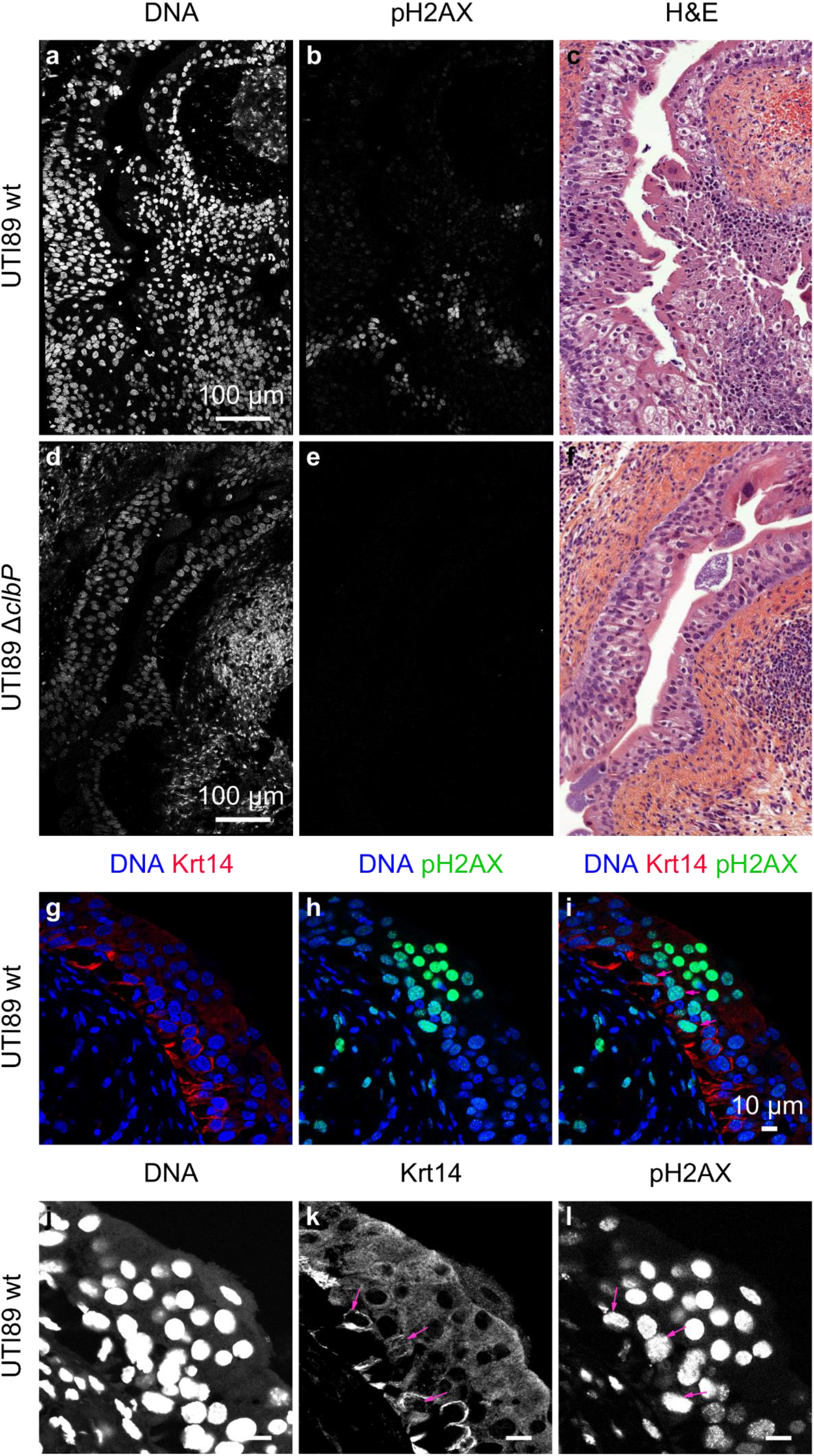
Colibactin induces DNA damage in urothelial cells of the regenerative compartment. **a-f.** DAPI (DNA: a, d), pH2AX (b, e) and haematoxylin-eosin (H&E: c, f) staining of paraffin-embedded bladder sections 24 hours after UTI89 wild-type (a-c) or UTI89Δ*clbP* (d-f) infection. The individual fluorescence channel images are shown in grayscale. See also Fig. S3. **g-l.** Paraffin-embedded bladders sections, were immuno-stained for pH2AX and Krt14, 24 hours post infection with wild-type UTI89. g-i: merged images: blue=DNA, red=Krt14, green=pH2AX. j-l: individual channel images shown at higher magnification in grayscale. Pink arrows identify cells positive for both Krt14 and pH2AX. Scale bar = 10 μm.

## Discussion

It is now increasingly clear that *pks*+ *E. coli* strains found in the intestinal microbiota may play a role in the aetiology and pathogenesis of colorectal cancer. These *E. coli* strains of phylogenetic group B2 are often the same ones responsible for UTIs, but to date, no study has been conducted on colibactin and UTIs. To the best of our knowledge the current study is the first to report the presence of a C14-Asn signature, a metabolite of colibactin production, in the urine of patients infected with *pks*+ UPEC. Thus, colibactin is produced during UTIs, one of the most common human bacterial infections worldwide. We used a mouse model of human UTI, to demonstrate that colibactin is produced and that it induces DNA damage in urothelial cells, specifically in bladder Krt14+ progenitor cells.

Our study confirms the frequent occurrence of *pks*+ UPEC observed in Europe and the US, irrespective of infection severity ^10–13^. This high prevalence in human subjects is in contrast to the apparent lack of selective advantages associated with the production of colibactin in UTIs mouse models. Our results suggest that colibactin is not essential for the ability of UPEC to colonise the bladder or to form IBCs, whereas colibactin has been shown to play a role in the pathogenicity of *E. coli* in extraintestinal infections such as septicemia or meningitis ^27,28^. However, it should be noted that the mouse model used omits a key step in the pathogenesis of UTIs: the domination and emergence from the intestinal reservoir ^29^. *E. coli* strains carrying *pks* island are commonly observed among strains with a greater ability to persist in the intestinal microbiota ^3^. In addition to the potential role of colibactin in modulating the intestinal microbiota and promoting gut colonisation, enzymes encoded by *pks* island are essential for the synthesis of siderophores and siderophore-microcins ^30,31^. Siderophores are major determinants in the domination of the intestinal niche ^32^. They confer upon strains that produce them the ability to outcompete other bacteria for iron, a rare essential nutrient. Moreover, siderophore-microcins, antimicrobial peptides which target phylogenetically-linked enterobacteria help UPEC to implant and proliferate within their intestinal reservoir, a step which precedes UTI ^10^.

The role of *pks* island in UPEC cannot merely be reduced to a determinant of intestinal colonisation. Indeed, our current study shows that the *pks* machinery is also active in the urinary tract and even within IBCs. Bacterial multiplication in these structures is intense, with a doubling time of nearly 30 min, which requires an efficient and optimised bacterial metabolism ^33^. Although colibactin is a small molecule, the metabolic cost of its production, *i.e.* expressing and operating the PK-NRP biosynthesis machinery is very high: nearly 1000 times higher than that of peptide synthesis ^34^. If UPEC trigger such energetically inexpedient assembly lines during UTIs, they must derive an adaptive benefit from it. One may speculate that other products of the machinery may confer this advantage. *In vitro*, there is indeed a wide diversity of metabolites produced by the *pks* machinery, ranging from other putative forms of “colibactins” (macrocyclic metabolites for example) to other smaller metabolites, which potentially vary between strains ^15,35^. The biological function of these metabolites remains to be elucidated, but some of these metabolites may be relevant to the pathogenesis of infections. We have, for instance, recently described the synthesis of C12-Asn-GABA, by the *pks* machinery of the probiotic Nissle 1917 strain and shown its digestive pain-relieving activity ^36^. The production of such an analgesic metabolite coupled with the increased production of siderophores by *pks*+ *E. coli* strains may provide a selective advantage to colonising the urinary tract, in addition to the digestive tract.

Irrespective of the role of *pks* island in the virulence of UPECs, the fact remains that these strains are genotoxic to bladder cells, particularly to bladder progenitor cells. Colibactin-induced DNA damage is incompletely repaired, resulting in the accumulation of gene mutations and cell transformation ^21,26,37^. We have previously shown that post-exposure chromosomal instability to colibactin can persist in daughter cells ^21^. DNA damage could also be propagated to contiguous cells by the induction of a senescence-associated secretory phenotype in the affected cells and the production of ROS, that could also explain the pattern of pH2AX positive patches of cells 24 hours after infection ^37,38^. In the current study, we observed that some murine bladder progenitor cells (Krt14+) display pH2AX positive nuclei. In humans, Krt14 expression characterises undifferentiated bladder cancers with particularly poor prognoses ^39,40^. Rodent bladder cancer models indicate that a significant proportion of cancerous tissue derives from Krt14 positive cells ^41^. Thus, colibactin damage to Krt14+ cells may initiate the formation and perpetuate the propagation of DNA lesions. The colibactin mutational signature has recently been identified in colorectal cancers but also in urinary tract cancers ^26,42^. Currently the main risk factors for bladder cancers are tobacco and occupational exposure to solvents, which are more frequently investigated than UTIs ^43^. However, a large worldwide bladder cancer case control study recently showed that regular UTIs were epidemiologically associated with an increased risk of urinary bladder cancer ^44^. Our findings suggest that *pks*+ UPEC UTIs may be an additional risk factor, particularly in cases of chronic and regular infections, irrespective of whether symptoms are present or not. We detected C14-Asn in the urine of patients with asymptomatic bacteriuria, which are usually not treated with antibiotics and can thus persist for periods of up to many years ^45^. A better understanding of the consequences of colibactin production could prompt a systematic search for *pks* island in UPEC isolates or C14-Asn in the urine of patients at risk.

## Materials and methods

### Bacterial strains

The archetypal *E. coli* strains used in this study were the UPEC strain UTI89 ^46^ and the colitogenic *E. coli* strain NC101 ^22^. The UTI89 and NC101 Δ*clbP* mutants were constructed using the lambda Red recombinase method ^47^ with primers IHAPJPN29 and IHAPJPN30, as previously described ^14^. The pMB702 construct (referred to as p-*clbP*) was used for complementation of the UTI89 Δ*clbP* mutant ^48^. For the *in vivo* complementation assay, the UTI89 Δ*clbP* mutant was transformed with the pCM17-*clbP* plasmid. Briefly, the *clbP* gene was PCR-amplified from pBRSK-*clbP* ^48^, with primers clbP-F-Bam_pm (5’-atGGATCCatgacaataatggaacacgttagc-3’) and pBRSK-F-Bam_pm (5’-atGGATCCcaagctcgga attaaccctc-3’) and cloned into the pCM17 vector ^49^ BamHI site. The ClbI C-terminal GFP fusion was constructed using the Gibson Assembly kit (New England Biolabs, MA, USA). Briefly, the *clbI* and *gfp* genes were amplified by PCR using the following primers: pK184-clbI-gb1(5’-GATTACGAATTCGAGCTCGGTACCCATGGCAGAGAATGATTTTGG-3’); clbI-gfp-gb2 (5’-CTTCTCCTTTTCCGCCTCCTCCGCCCTCATTAATCATGTCGTTAACTAG-3’); clbI-gfp-gb3 (5’-GATTAATGAGGGCGGAGGAGGCGGAAAAGGAGAAGAACTTTTCACTGG-3’) and gfp-pK184-gb4 (5’-TGCAGGTCGACCTCGAGGGATCCCCTTATTTGTATAGTTCATCCATGCC-3’). A glycine linker (5’-GGCGGAGGAGGCGGA-3’) was introduced at the *clbI* and *gfp* junction for flexibility. PCR amplified fragments and SmaI-digested pK184 vector were assembled according to the manufacturer’s recommendations.

### Collection of clinical strains and urines

The collection of 225 *E. coli* strains from urine samples at the Adult Emergency Department of Toulouse University Hospital, France, between July and October 2017, was previously described ^10^. According to the French regulations relating to observational database analyses, the study did not require specific informed consent. Urine was collected from 223 patients (women and men, under 75 years) with either pyelonephritis (104), symptomatic infections excluding pyelonephritis (cystitis) (83), or asymptomatic bacteriuria (36) without urological comorbidities or catheterisation. All strains were identified as *E. coli* by matrix-assisted laser desorption/ionisation time-of-flight mass spectrometry (Microflex LT MALDI-TOF MS, Bruker Daltonik GmbH, Germany). Samples of the corresponding urine collected in boric acid tubes were stored at −80 °C until lipid analysis.

### C14-Asn quantification

Extraction and quantification of C14-Asn was performed as previously described ^36^. Briefly, 5 μL internal standard mixture (Deuterium-labelled compound at 400 ng/mL) was added to the bacterial pellets of 24 h DMEM cultures or to 500 μL of urine samples before crushing, followed by addition of cold methanol (MeOH) (15% final volume) and homogenisation. After centrifugation, supernatants were solid phase extracted on HLB plates (OASIS® HLB 2 mg, 96-well plate, Waters, Ireland). Following washing with H2O/ MeOH (90:10, v/v) and elution with MeOH, samples were evaporated twice under N2 and finally resuspended in 10 μL MeOH. Separation and quantification of C14-Asn was performed on a high-performance liquid chromatography coupled to tandem mass spectrometry system (G6460 Agilent) ^36^. Limit of detection was 2.5 pg and limit of quantification was 5 pg.

### Sequencing data, sequence alignments and phylogenetic analyses

Whole genome sequencing was performed using the Illumina NextSeq500 Mid Output platform (Integragen, Evry, France) to generate 2 × 150 bp paired-end reads, at approximately 80x average coverage. Genome *de novo* assembly and analysis were performed with the BioNumerics 7.6 software (Applied Maths) and Enterobase (http://enterobase.warwick.ac.uk/). For SNP-based phylogenetic trees, core genome alignments were generated after mapping raw reads to the *E. coli* MG1655 genome. The core genome phylogenetic tree was inferred with the Maximum-likelihood algorithm using Enterobase for B2 phylogroup strains.

### Exogenous DNA cross-linking assay

As previously described ^18^, the pUC19 plasmid, linearised with BamHI, was exposed to bacteria pre-grown (3 × 10^6^ CFU) in DMEM 25 mM Hepes for 40 min at 37 °C. The DNA was purified and 100 ng submitted to denaturing gel DNA electrophoresis (40 mM NaOH and 1 mM EDTA, pH ~12.0). After a neutralisation step, the gel was stained with GelRed and visualised under UV using the ChemiDoc Imaging System (BioRad).

### pH2AX and pRPA immunofluorescence analysis of post infected HeLa cells

HeLa cells (ATCC CCL2) were cultured and infected in 8-well chamber slides (Labtek), as previously described ^18^. 24 hours after passaging, Hela cells were infected with bacteria precultured in DMEM 25 mM HEPES at a multiplicity of infection (MOI) of 50, for 4 h. Cells were then washed and incubated overnight in cell culture medium with 100 μg/ml gentamicin. Immunofluorescence analysis was then performed as previously described ^18^. Briefly, cells were pre-extracted in PBS 0.1%, Triton X-100 and fixed in PBS 4% formaldehyde, permeabilised, blocked with MAXblock medium (Active Motif) and stained with antibodies against pH2AX (1:500, JBW301, Millipore) and S33p-RPA32 (1:500, A300-264A, Bethyl), diluted in MAXblock 0.05%, Triton X-100. Cells were washed 3 times for 5 min in PBS 0.05%, Triton X-100 and incubated with secondary AlexaFluor 488 or 568 (Invitrogen) diluted 1:500 in MAXblock medium with 1 μg/ml DAPI (Sigma). Slides were mounted with Fluoroshield (Sigma) and examined on a Leica SP8 laser scanning confocal microscope in sequential mode with LasX software (Leica), while keeping the same laser and detector settings between different wells. Final images were processed and edited with the ImageJ software.

### Mouse UTI model

Animal infections were performed in accordance with the European directives for the protection of animals used for scientific purposes (2010/63/EU). An ethics committee approved the protocol (number CEEA-122 2014-53). Female, 6-8 weeks old, C3H/HeN mice (Janvier Labs) were infected transurethrally as previously described ^50^. Briefly, UPEC were cultivated statically in LB ^50^ and resuspended to an inoculum of 10^8^ CFU in 50 μl PBS. Mice under 4% isoflurane anaesthesia were inoculated twice at one-hour intervals with a pump that delivered 10 μL/s. At 6 or 24 hours, mice were euthanised, bladders were harvested, homogenised for bacterial enumeration on agar plates, stored in OCT compound at −80 °C or fixed in 4% formaldehyde after filling the bladder prior to paraffin embedding. Each experiment was conducted in duplicate with 5 to 8 mice per group.

### Histological and immunofluorescence analyses of mouse bladder tissue

Histological bladder slices were prepared using standard protocols and processed for haematoxylin-eosin staining then scanned using a Pannoramic 250 scanner (3DHistech). Final images were captured using CaseViewer software (3DHistech).

For GFP and pH2AX detection on OCT compound-embedded bladders, 8 μm thick sections were dried on Superfrost plus glass slides, fixed for 15 min in 4% formaldehyde then rinsed with PBS, blocked and permeabilised with MaxBlock, 0.3% Triton X100. Primary anti-GFP (Rabbit, Abcam 6556) and anti-pH2AX (Mouse, Millipore #O5-636) antibodies were diluted at 1:200 in MaxBlock, 0.3% triton X100. After washing in PBS, 0.05% Triton X100, slides were incubated with secondary antibodies at 1:200 (Alexa 633 Goat anti-Rabbit antibody (Invitrogen A21071) and Alexa 488 Goat anti-Mouse antibody (Life technologies A11029)) and DAPI. For pH2AX and Krt14 staining on paraffin-embedded bladders, 5 μm thick sections were de-paraffinised and re-hydrated in xylene, ethanol and tap water. Unmasking was performed with a 6-min trypsin digestion (Trypsin-EDTA solution, Sigma, T392) followed by 30 min incubation in citrate buffer pH 6 at 80–95°C. Sections were then blocked and permeabilised in MaxBlock, 0.3% Triton X100. Slices were stained with anti-pH2AX (Rabbit, Cell Signalling S9718) at a dilution of 1:200 and anti-Krt14 antibodies (Chicken, Ozyme BLE906001) at a dilution of 1:250 in MaxBlock, 0.3% Triton X100. Sections were washed in PBS, 0.05% Triton X100 and incubated with secondary antibodies (Alexa 633 Goat anti-Rabbit antibody, Invitrogen A21071, at a dilution of 1:200; Alexa 555 Goat anti-Chicken antibody, Invitrogen A32932, at a dilution of 1:500). Images were acquired as with infected cultured cells.

### Fluorescence *in situ* hybridisation (FISH) of mouse bladder tissue

Five micron paraffin-embedded bladder sections were de-paraffinised in xylene/ethanol. Sections were incubated in lysozyme solution (10 mg/ml, Sigma, France) for 15 minutes at 37 °C and exposed to 100 μl of universal bacterial 16 S fluorescent rRNA probe (Eub338, GCTGCCTCCCGTAGGAGT-Cy5’, Eurofins, France) at a concentration of 5 ng/μl, in hybridisation buffer (20 mM Tris-HCl, pH7.4, 0.9 M NaCl, 0.01% SDS) at 46°C for 3 hours ^51^. Sections were then incubated in a 48 °C prewarmed saline-sodium citrate wash buffer (30 mM sodium citrate, 300 mM sodium chloride, pH7.4, Invitrogen, France) for 20 minutes. To stain polysaccharide-rich content, sections were counterstained with FITC-conjugated wheat germ agglutinin (Invitrogen, France) at a 1:1000 dilution in PBS buffer for 30 min. Slides were mounted with Fluoroshield containing DAPI. Images were acquired using a confocal laser scanning microscope (Zeiss LSM 710) and final images were processed and edited with the ImageJ software.

### Statistical analyses

Graphical representation and statistical analyses were carried out using GraphPad Prism 8.3. P values were calculated using two-tailed Mann-Whitney U test.

## Acknowledgements

The authors thank Alexandre Perrat, Frédéric Auvray and Laurent Cavalié for their help with genomic DNA extraction and sequence analysis and Pauline Dragot for her excellent technical help. We thank the technical microbiology laboratory staff from the Toulouse University Hospital for helping to culture and isolate *E. coli* strains. The authors also wish to thank the staff of the Tri GenoToul imaging facility, Toulouse (particularly Sophie Allart and Danièle Daviaud), the experimental histology facility (CREFRE, Toulouse) and the lipidomic facility of MetaToul, Toulouse. The authors thank the team members for their helpful discussions and critical reading of the manuscript and Sébastien Déjean for his help with data analysis.

This work was supported by grants from the French National Agency for Research (ANR) (UTI-TOUL ANR-17-CE35-0010 and LiBacPain ANR-18-CE14-0039). C.V.C was supported by a two-year grant from the French National Institute for Health and Medical Research (poste d’accueil INSERM 2018). The funding bodies did not contribute to the study design, collection of data, interpretation of results or to the decision to submit the work for publication.

## Authors contribution

C.M., C.V.C. collected human clinical samples (urines and strains). C.M., C.V.C, E.O. analysed the strains. C.V.C., P.L.F., N.C. performed lipid extractions and analyses. A.S., N.B.G., P.M., C.V.C. designed and constructed the molecular engineered strains. C.V.C., N.B.G., J-P.N. performed *in vitro* genotoxicity assays. C.F., M.F., P.M. designed and carried out *in vivo* experiments. P.M., C.B. and C.V.C. analysed *in vivo* samples. C.V.C., J-P.N., J-P.M. performed histological and microscopic analyses. C.V.C., C.M., J-P.N., J-P.M., P.M., N.C., M-P.R., H.C., M.F., E.O. contributed to the analysis and interpretation of the data. C.V.C, J-P.N., C.M., E.O. drafted and wrote the paper. E.O. obtained dedicated funding to support this work.

