## Supplementary data for "Uropathogenic *E. coli* induces DNA damage in the bladder"

**Table S1: Human *pks*+ UPEC belong to major lineages of phylogroup B2 ExPEC.** Phylogroup, Sequence Types and presence of a *pks* island were determined from whole genome sequences of UPEC isolates, along with the detection of C14-Asn from the corresponding urine.

| Phylogroup | **B2** | | | | | | | **D** | | | **Others** | **Tot.** |
|---|---|---|---|---|---|---|---|---|---|---|---|---|
| Sequence Type | 73 | 95 | 141 | 404 | 131 | Oth. | Tot. | 69 | Oth. | Tot. | Others | |
| n | 29 | 27 | 16 | 14 | 23 | 46 | **155** | 26 | 8 | **34** | **36** | 225 |
| Infection | | | | | | | | | | | | |
| Asymptomatic bacteriuria | 4 | 3 | 6 | 0 | 4 | 10 | 27 | 3 | 2 | 5 | 5 | 37 |
| Cystitis | 13 | 2 | 6 | 8 | 13 | 19 | 61 | 6 | 1 | 7 | 16 | 84 |
| Pyelonephritis | 12 | 22 | 4 | 6 | 6 | 17 | 67 | 17 | 5 | 22 | 15 | 104 |
| *pks* island | | | | | | | | | | | | |
| *pks*+ | 29 | 7 | 16 | 14 | 0 | 30 | 96 | 0 | 0 | 0 | 0 | 96 |
| C14-Asn+ | 13 | 6 | 12 | 5 | 0 | 19 | 55 | 0 | 0 | 0 | 0 | 55 |

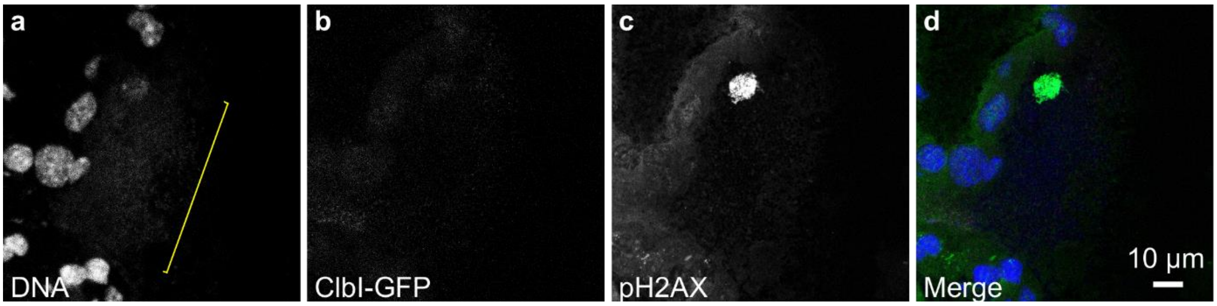


**Fig. S1: Immunofluorescence staining of GFP and pH2AX on frozen bladder sections 6 hours post infection with wild-type UTI89, non-expressing GFP.** The individual channel images are shown in grayscale. In the merge (d) image: blue=DNA, magenta=ClbI-GFP, green=pH2AX, grey=phase contrast. The yellow bracket shows an umbrella cell containing an IBC.

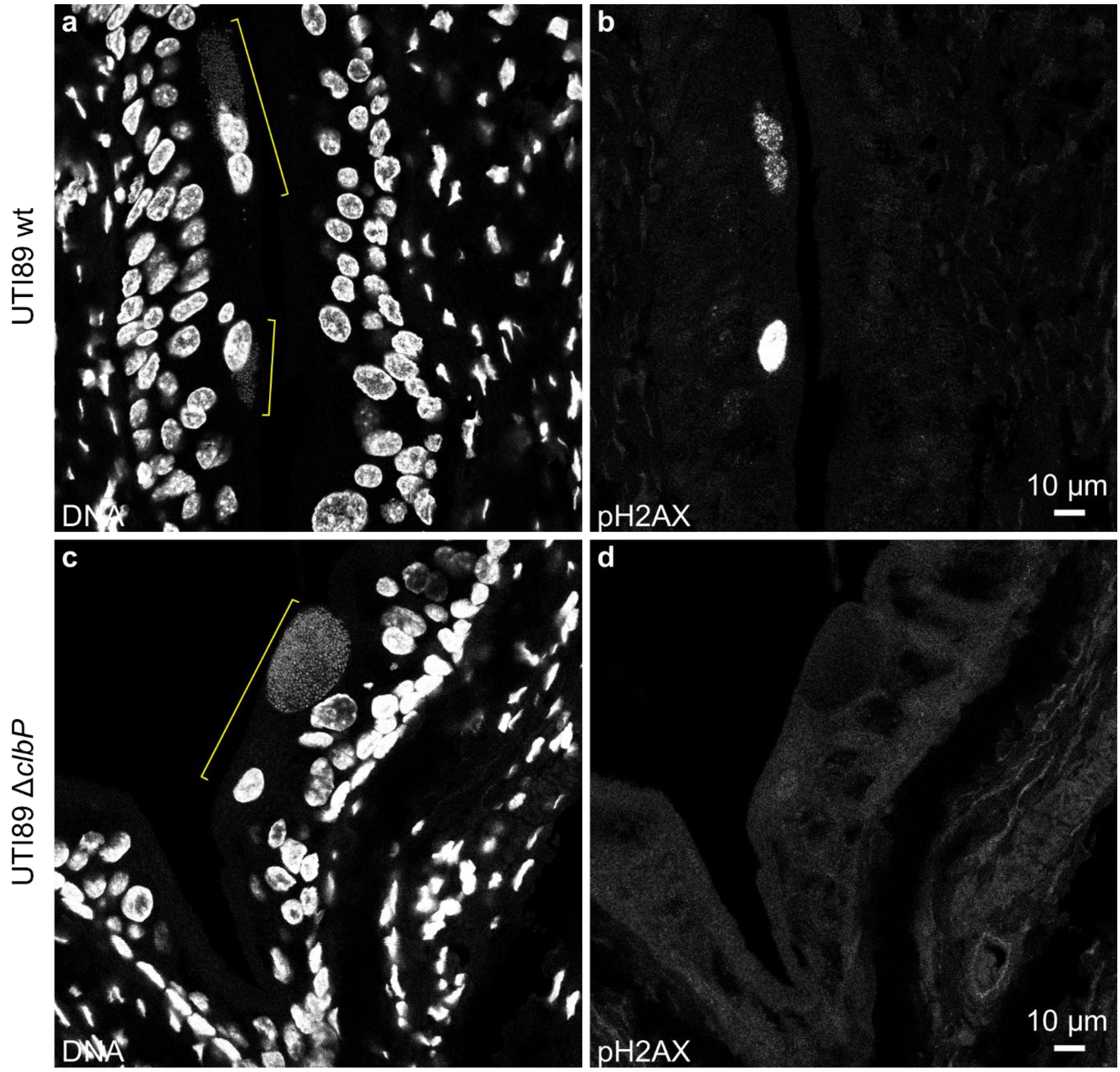


**Fig. S2: Colibactin induces DNA damage in umbrella cells containing IBCs at early stages of infection**. Immunofluorescence staining of pH2AX on paraffin embedded bladders sections 6 hours post-infection with the wild-type UTI89 (a, b) or the Δ*clbP* isogenic mutant (c,d). The individual channel images are shown in grayscale. Yellow brackets show umbrella cells containing IBCs, with DAPI stained bacterial colonies.

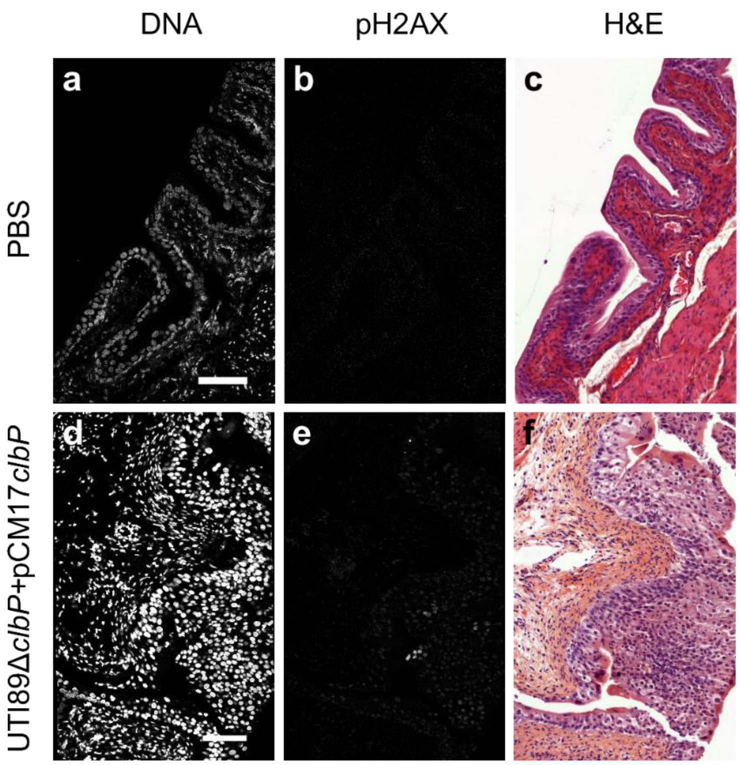


**Fig. S3: Colibactin induces DNA damage in urothelial cells. a-f.** Haematoxylin-eosin (H&E: c, f), DAPI (DNA: a, d) and immunofluorescence staining of pH2AX (b, e) on paraffin-embedded bladders sections 24 hours post PBS inoculation (a-c) or infection by UTI89Δ*clbP*+pCM17*clbP* (d-f). For immunofluorescence, the individual channel images are shown in grayscale. Scale bar = 100 µm.

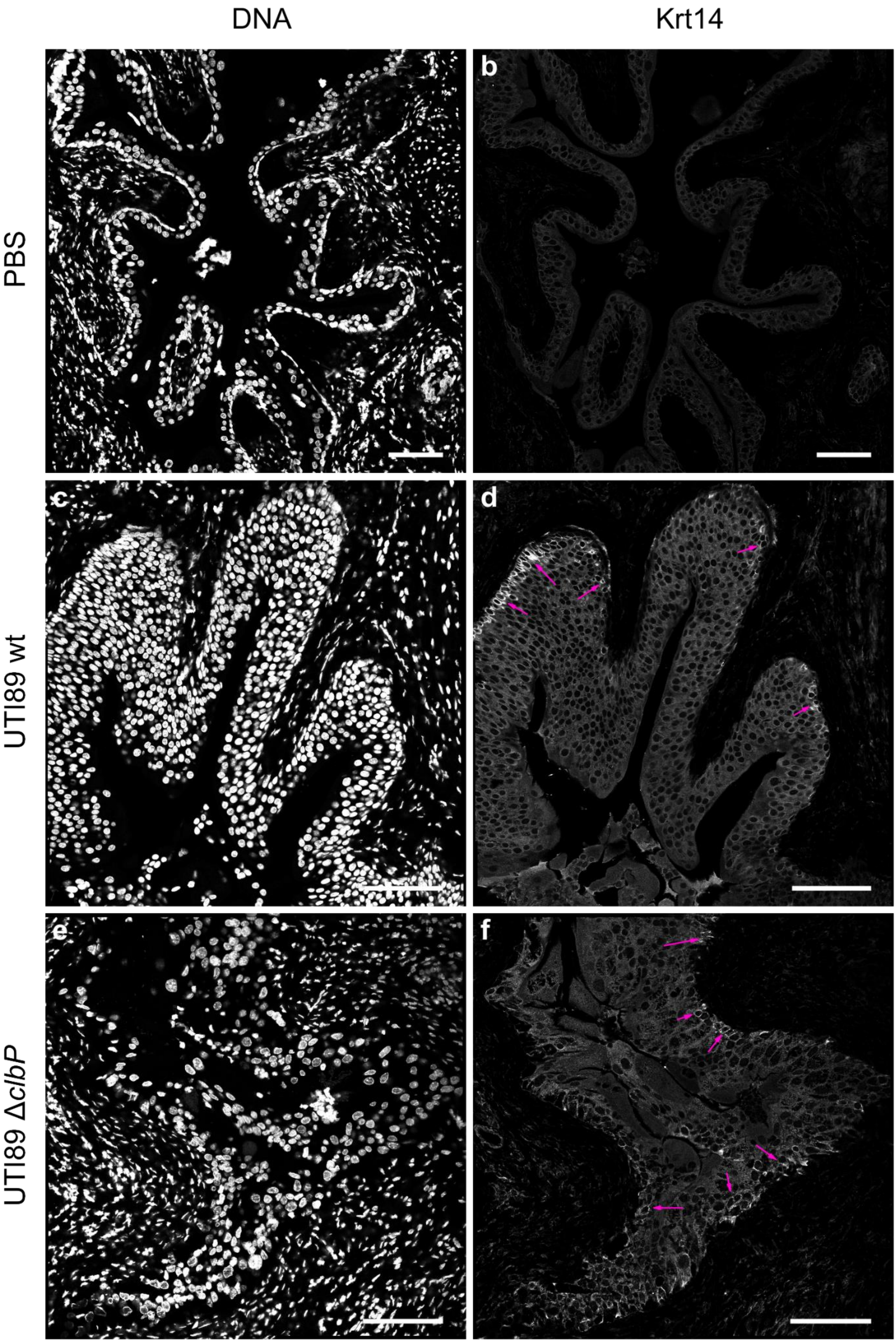

DNA
Krt14
PBS
UTI89 wt
UTI89 ΔclbP
b
c
d
e
f

**Fig. S4: Krt14 positive cell proliferation is induced upon infection.** Immunofluorescence staining of Krt14 and DAPI stained DNA on paraffin-embedded bladders sections 24 hours post PBS inoculation (a-b) or infection by UTI89 wild-type (c-d) or UTI89 Δ*clbP* (e-f). The individual channel images are shown in grayscale. Scale bar = 100 µm. Pink arrows: Areas with Krt14+ cells after infection.
